## Supplement DM Tutorial for "Skeletonization of neuronal processes using Discrete Morse techniques from computational topology"

### Supplementary Materials

#### S.1 DM Tutorial

##### S.1.1 Morse theory and 1-stable manifolds in the smooth case

We first provide an informal description of the relevant part of Morse theory that motivates the graph reconstruction algorithm we use. Let $f : \mathbb{R}^d \to \mathbb{R}$ be a smooth function on $d$-dimensional Euclidean space. In our applications for processing neuron images, the domain is either $\mathbb{R}^2$ for 2D images or $\mathbb{R}^3$ for 3D volumetric image data. The gradient vector at a point $\mathrm{p} \in \mathbb{R}^d$ is defined as [1]:

$$\nabla f(\mathrm{p}) = -[\frac{\partial f}{\partial \mathsf{x}_1}, \frac{\partial f}{\partial \mathsf{x}_2}, \ldots, \frac{\partial f}{\partial \mathsf{x}_d}]^T,$$

where $(\mathsf{x}_1, \ldots, \mathsf{x}_d)$ represents an orthonormal coordinate system for $\mathbb{R}^d$. In simple terms, $\nabla f(\mathrm{p})$ represents the *steepest descending direction* along which the function $f$ decreases fastest when moving away from $x$, and its norm $\|\nabla f(\mathrm{p})\|$ is the rate of this change. Gradient vectors for all points in $\mathbb{R}^d$ form a vector field on $\mathbb{R}^d$, called the *gradient vector field*. A point p with vanishing gradient; that is, $\nabla f(\mathrm{p}) = [0, 0, \ldots, 0]^T$, is called a *critical point*; otherwise, it is a regular point. A critical point p of $f$ is *non-degenerate* if the Hessian matrix (formed by all second-order partial derivatives $[\frac{\partial^2 f}{\partial \mathsf{x}_i \partial \mathsf{x}_j}]$) has full rank; otherwise, it is degenerate.

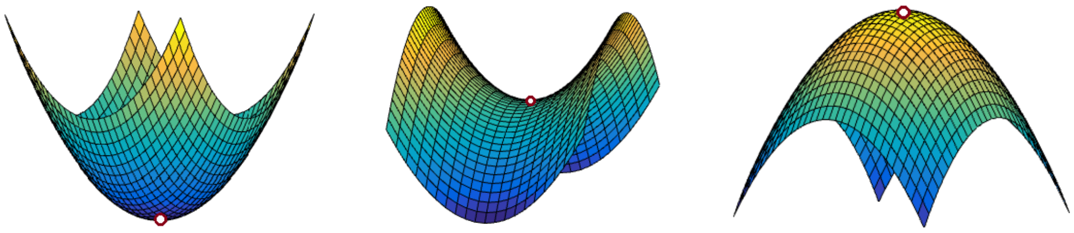

min (index 0) saddle (index 1) max (index 2)

**Supplementary Fig. 1** | Three types of critical points (of index 0, 1, and 2) for a Morse function defined on $\mathbb{R}^2$.

A *Morse function* is a smooth function where all critical points are (i) non-degenerate and (ii) have distinct function values. Morse functions are well-behaved functions whose critical points also have nice characterizations. For example, by Morse Lemma [46], for a Morse function $f$ on $\mathbb{R}^d$, there are $d + 1$ types of critical points, local minima (index 0), local maxima (index $d$), and $(d - 1)$-types saddle points (of indices from 1 to $d - 1$). In the case of a 2D Morse function $f : \mathbb{R}^2 \to \mathbb{R}$, there are three types critical points, local minima, local maxima, and saddle points as shown in Fig. 1. Critical points capture local behavior of the function $f$. The global variation can be partially captured by concepts such as integral lines and (un)stable manifolds. In particular, an *integral line* of $f$ is a maximal path in $\mathbb{R}^d$ such that the tangent vector of this path at any point coincides with the gradient $\nabla f(x)$. Intuitively, imagine that we view the graph of the function $f : \mathbb{R}^d \to \mathbb{R}$ as a "terrain" defined in $\mathbb{R}^{d+1}$; see Fig. 2 for an illustration where the last coordinate (i.e, the height of each point) corresponds to $f(x)$ at each $x \in \mathbb{R}^d$. The lift of an integral line to the surface of terrain is a "flow line" on the terrain that a water drop will follow when flowing in the direction of the steepest descending direction at any point. See the green solid curve in Fig. 2 for an example of a flow line, which starts at maximum $t_1$ and ends at minimum $v$.

Consider a flow line. The water drop will keep flowing till it reaches a point where there is no descending direction – these are exactly points where gradients vanishes, namely critical points. Hence flow lines (thus integral lines) have to "start" and "end" at critical points. The *unstable manifold* of a critical point is the union of points along all integral lines that ultimately "end" at this point. We are particularly interested in the 1-unstable manifolds, which are those flow lines that end at index $d - 1$ saddle points. In the example of a function defined in $\mathbb{R}^2$, these are pieces of curves that connecting maxima with saddles; see the white dotted curves in Fig. 2.

Such 1-unstable manifolds for saddles may be conceptualized of as "mountain ridges" of this terrain (graph of function $f$), connecting mountain peaks (maxima) to peaks (maxima) via saddles, and separating different valleys (basins of minima). There is also a dual concept of 1-*stable manifolds*, that correspond to "valley ridges", connecting minima to minima via saddles and separating mountain peaks.

[1] Note that this is a negative version of the sign convention used in classical Morse theory. We use this negated version as it is then aligned with the steepest descending direction of $f$; while the usual notion of the gradient vector indicates the steepest ascending direction.

In our setting, we can view a 2D image or a 3D image as a real valued density field on $\mathbb{R}^2$ or $\mathbb{R}^d$, respectively. Consider the terrain (graph) of this density field. The mountain ridges of this terrain can capture where strong signals of cell processes are. We will aim to use the 1-unstable manifold of this density function to capture its mountain ridges and further to capture the neuronal cell processes.

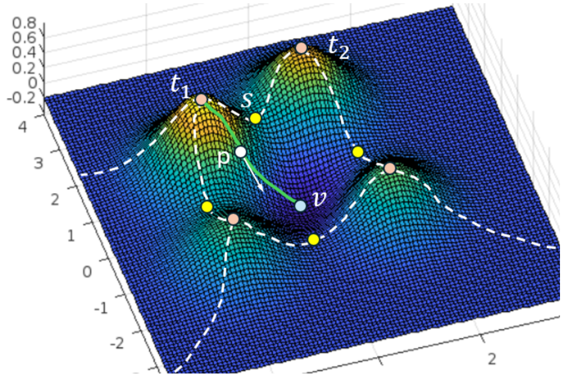


**Supplementary Fig. 2 |** White dotted lines are the union of 1-unstable manifolds. The green curve is an example of a flow line. Pink points are maxima, yellow points are minima, while the blue point $v$ is a minimum.

#### S.1.2 Discrete Morse theory

Numerical data on a computer are discretely sampled and do not equal the mathematical notion of a smooth function. We can view the sampled 2D or 3D image data that as a discretization of a smooth function $\rho$ defined on $\mathbb{R}^2$ or $\mathbb{R}^3$. While one could compute the 1-unstable manifolds from a continuous extension of the discrete evaluations of $\rho$ at pixels (e.g, a piecewise-linear approximation), this could to be sensitive to approximation and numerical error as (un)stable manifolds are defined based on gradients. Simplifying / denoising the resulting (un)stable manifolds could be challenging.

Discrete Morse theory, a combinatorial analog of the classical Morse theory for cell complexes, was proposed by Forman[23, 47] as a mathematically well defined but explicitly discrete and computationally viable approach to Morse theory suitable for analysis of real life data sets.

We provide a brief informal description of some concepts from discrete Morse theory that are relevant to the present DM-based graph reconstruction algorithm. We utilize simplicial complexes, which are complexes consisting of building blocks called simplices, which in 2D consist of points, line segments, triangles and pyramids, glued appropriately along their faces. A $d$-dimensional (geometric) simplex is the convex combination of $d+1$ affinely independent points. Thus 0, 1, and 2-simplices correspond to vertices, edges, and triangles. See Fig. 3 for an example of a 2-dimensional simplicial complex, triangulating a square, consisting of a collection of vertices, edges, and triangles. In our applications, we use cubical complexes instead of simplicial complexes to represent images, where pixels are vertices, and cells are squares instead of triangles. However, we will simplicial complexes to illustrate these concepts for simplicity of presentation.

Given a simplicial complex $K$, a *discrete gradient vector* is a pair of simplices $(\sigma, \tau)$ such that $\sigma$ is a co-dimension one face of $\tau$; e.g., $\tau$ is an edge incident on a vertex $\sigma$, or $\tau$ is a triangle incident on an edge $\sigma$, and so on. Note that a discrete gradient vector is therefore a combinatorial pair, instead of a true vector. Nevertheless, a given pair, say $(\sigma, \tau)$, still indicates a "flow direction" from $\sigma$ to $\tau$, much like in the case of smooth Morse theory; see Fig. 3 where each discrete gradient vector $(\sigma, \tau)$ is indicated by a vector from $\sigma$ to $\tau$. A *V-path* is a sequence of simplices $\sigma_0, \tau_0, \sigma_1, \tau_1, \ldots, \sigma_k, \tau_k, \sigma_{k+1}$ such that for any $i \in [0, k]$, we have (i) the pair $(\sigma_i, \tau_i)$ is a discrete gradient vector, and (ii) $\sigma_{i+1}$ is a co-dimension one face of $\tau$. A V-path is *cyclic* if $\alpha_0 = \alpha_{k+1}$; otherwise, it is *acyclic*. See Fig. 3 for some examples.

Given a discrete gradient vector field $M$ of $K$, a simplex $\sigma$ is *critical* if it is not involved in any pair (ie., discrete gradient vector) in $M$; intuitively meaning that the gradient is "vanished" at $\sigma$. The index of this critical simplex is its dimension. Intuitively, given a triangulation $K$ of a 2D domain and a discrete gradient vector field $M$, critical vertices, critical edges, and critical triangles are analogous to minima (index-0 critical points), saddle (index-1) and maxima (index-2) for a smooth Morse function defined on this domain. The 1-unstable manifolds corresponding to saddles are V-paths connecting critical edges (saddle) to critical triangles (maxima). Each such V-path is a sequence of alternating edges and triangles. If $K$ is a triangulation of a $d$-dimensinal domain, such a V-path will be sequence of $(d-1)$-simplices and $d$-simplices, which can be expensive to compute and manipulate. Hence in practice, following [22, 24], we use the following trick: Instead of aiming to compute the 1-unstable manifold of an input function (viewed as a density field) $\rho$ to capture the mountain ridges, we calculate the *1-stable manifolds* of the negation of $\rho$, that is, for $-\rho$. The 1-stable manifolds capture "valley ridges" connecting the minima (bottom of the valleys) with saddles. In the discrete Morse setting, 1-stable manifolds correspond to V-paths connecting critical edges to critical vertices

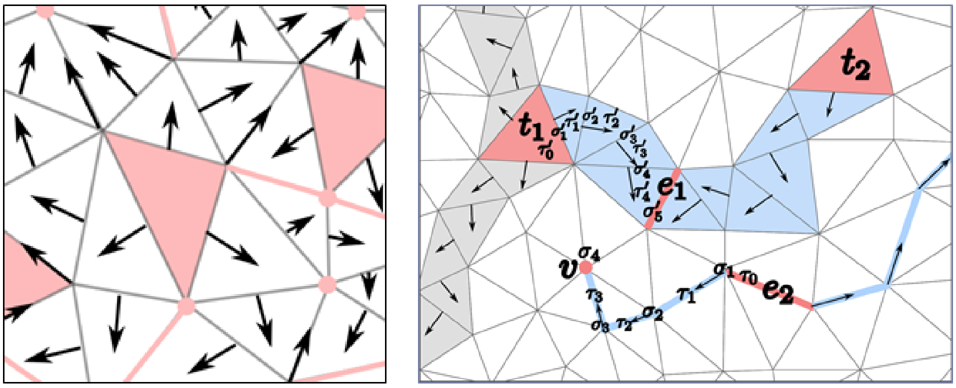


(a) (b)

**Supplementary Fig. 3 |** (a) A discrete gradient vector field. (b) Examples of V-paths analogous to 1-unstable manifolds (connecting $t_1/t_2$ to $e_1$) and 1-stable manifolds (connecting $v$ to $e_2$). (Image courtesy of [3].)

via sequence of alternating vertices and edges (regardless of the dimension of the domain), which are much easier to compute and maintain.

Finally, another advantage of using discrete Morse theory is that it provides a simple and combinatorial method for simplifying a given discrete gradient vector field $W$ to $W'$ to reduce the number of critical simplices. This is achieved via the so-called *Morse cancellation* operation: Specifically, a pair of critical simplices $\alpha$ and $\beta$ of dimension $p$ and $p+1$ (e.g, a critical vertex $\sigma$ and a critical edge $\beta$) is *cancellable* if there is a **unique** V-path connecting them. Given such a cancellable pair of critical pairs $(\alpha, \beta)$, we can simply "invert" the direction of the gradient vectors in this path and this will produce a new gradient field where $\alpha$ and $\beta$ are no longer critical, while other critical simplices stay the same. This operation can be repeatedly performed as long as there are cancellable pairs of critical simplices to reduce the number of critical simplices, and thus simplify the discrete gradient vector field.

Note that any pair of critical simplices can be cancelled as long as the unique V-path condition is satisfied. To decide which pairs to simplify so as to remove the "noise" in the input density field, we will use another topological concept called the persistent homology which we briefly introduce next.

##### S.1.3 Persistent homology

Persistent homology [3–5, 48–50] is a fundamental recent development in the field of computational topology, underlying many topological data analysis methods. Below we provide an intuitive description about it, to help explain how it can be used to measure importance of critical points. See [3] for more detailed introduction of these topics.

First we describe the idea for the smooth setting, where we assume that we are given a smooth function $f : X \to \mathbb{R}$ defined on a space $X$ (e.g, $X$ is a square for the case of 2D image). Now imagine we trace how the function $f$ evoles on $X$ via the following growing sequence of subspaces of $X$:

$$X_{t_1} \subseteq X_{t_2} \subseteq \cdots \subseteq X_{t_m} = X, \text{with} \;\; t_1 < t_2 < \cdots t_m,$$

where $X_t := \{x \in X \mid f(x) \leq t\}$ is the so-called *sub-level set of $f$ at $t$*. This is called the *sub-level set filtration of $X$ w.r.t. $f$*, which intuitively sweeps $X$ by increasing $f$ values, and tracks the subspace $X_t$ already swept. In particular, during this process, sometimes *new topological features* (such as components, loops/handles, voids) can appear, and sometimes they disappear. These topological features can be captured algebraically by the so-called homology classes. The creation (birth) and destruction (death) of such features can be captured by the so-called *persistent homology* [49], which can be summarized by a simple summary, called the *persistence diagram*. More precisely, it turns out that the birth and death of topological features (homology classes) can only happen when the sublevel set $X_t$ sweeps through a critical point of $f$. We can therefore track the birth and death events by a collection of *persistent pairings* $\Pi_f = \{(v_b, v_d)\}$, where each pair $(v_b, v_d)$ contains the critical points where certain topological feature is created and killed. Their function values $f(v_b)$ and $f(v_d)$ are referred to as the *birth time* and *death time* of this feature. The corresponding collection of pairs of (birth-time, death-time) is called the *persistence diagram*, formally, $\mathrm{dgm}(f) = \{(f(v_b), f(v_d)) \mid (v_b, v_d) \in \Pi_f\}$. Each *persistent point* $(f(v_b)), f(v_d))$, corresponding to the birth time and death time of some homological features through the filtration, gives rise to a point in the *birth-death plane*, as shown in Fig. 4 where we provide a simple example of 1D function. (For example, in this example, as we sweep pass minimum $x_3$, a new component is created in the sub-level set. This component is merged to an older component (created at $x_1$) when we sweep past critical point (maximum) $x_4$. This gives rise to a persistence pairing $(x_3, x_4)$ corresponding to the point $(f_3, f_4)$ in the persistence diagram.) The importance of the topological feature captured by the persistent pairing $(v_b, v_d)$ is captured by its *persistence*, defined as $|f(v_d) - f(v_d)|$, as it measures the "lifetime" of this topological feature through the filtration.

Equivalently, one can also represent a persistent point $(t_b, t_d)$ as an interval, called a *persistent bar*, and the collection of persistent points then give rise to a collection of "bars", giving rise to a representation called *persistent barcode*. In Fig.2 of the main text, we used the persistence barcode representation to make the relation to the persistent pairing giving rise to each bar more clear.

Finally, we note that there is an algorithm [9] based on matrix reduction to compute persistent homology, persistent pairings and the resulting persistence diagram.

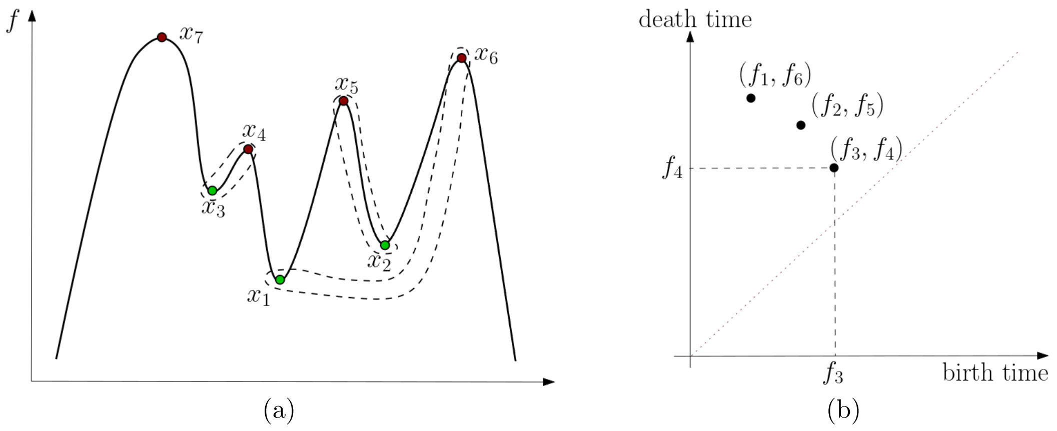


**Supplementary Fig. 4 |** (a) A simple 1D function $f : \mathbb{R} \to \mathbb{R}$. Its persistence pairings of critical points are marked by the dotted curves: $\Pi_f = \{(x_1, x_6), (x_2, x_5), (x_3, x_4), \ldots\}$. (b) shows its corresponding persistence diagram $\mathrm{dgm}(f) = \{(f_1, f_6), (f_2, f_5), (f_3, f_4), \ldots\}$, where $f_i = f(x_i)$ for each $i \in [1, 6]$.

**Persistence algorithm in discrete setting.** In the discrete setting, suppose we are given a triangulation $K$ and a function $f : V \to \mathbb{R}$ defined at vertices of $K$. We can simulate the above sub-level set filtration by the so-called *lower-star filtration*. Specifically, one can think that the function $f$ on $V$ is extended to a function on all simplicies in $K$ by $f(\sigma) = \max_{v \in \sigma} f(v)$, for each simplex $\sigma \in K$ (e.g, for an edge $\sigma$, its function value $f(\sigma)$ equals the larger function value of its two vertices). One can then take the sublevel-set filtration of this simplex-wise valued function. Next, we can run the standard persistence algorithm [9] to this filtration, and the output is a collection of *pairs of simplices* $\Pi_f$. For each pair of simplices $(\sigma, \tau) \in \Pi_f$, it intuitively captures the birth and death of some homological feature in the sublevel sets. Analogous to the smooth setting, this pair of simplices gives rise to a persistent point $(f(\sigma), f(\tau))$ in the persistence diagram, and its persistence $per(\sigma, \tau) = |f(\tau) - f(\sigma)|$ measures the lifetime (importance) of this feature. We will use these pairs and their importance as captured by the persistence to guide the simplification of discrete gradient vector field (and thus the resulting 1-(un)stable manfiolds).

##### S.1.4 Persistence-guided discrete Morse based graph reconstruction algorithm

We now put all pieces together and introduce the persistence-guided discrete Morse based graph reconstruction algorithm [24, 44], denoted by `DiMorSC` ().

On the high level, given a density function $\rho : X \to \mathbb{R}$ on a domain $X \subset \mathbb{R}^d$, note that we will consider the negation $f = -\rho$ of the density function, and aim to compute the 1-stable manifolds (instead of 1-unstable manifolds) to capture the valley ridgets (instead of mountain ridges). This is for the purpose to simplify the manipulation of discrete gradient vector field in the discrete Morse setting. Intuitively, we first compute the persistence pairings of all critical points of the function $-\rho$. We will then simplify the density function $\rho$ by "canceling" those pairs of critical points with low persistence, and only consider the 1-stable manifolds of remainder saddles with large persistence bigger than a given threshold $\delta$. The union of such 1-stable manifolds will capture important valley ridges of $-\rho$ (and thus mountain ridges of the density map $\rho$), and is the output reconstructed graph.

The above intuition can be translated to the discrete Morse setting, and we present the resulting algorithm in Algorithm 1, which is based on the simplified algorithm proposed by [22].

The original algorithm takes a triangulation $\mathsf{K}$ of the domain of interests and a density function $\rho$ given at the vertices of $\mathsf{K}$ as input. In our case, since our inputs are 2D images, instead of a triangulation, we take $\mathsf{K}$ to be the 2-skeleton (vertices, edges, and squares) of the 2D-cubical complex of the domain and use PHAT to compute persistence directly for such cubic complex. This does not change any part of the algorithm. Note that the algorithm will take as input a user-defined persistence threshold $\tau$; only 1-stable manifolds of saddles (critical edges) with persistence larger than $\tau$ will be computed and output.

**Algorithm 1** $G$ = `DiMorSC` (K, $\rho$, $\tau$)

1: Persistence Computation
  - Compute persistence pairings induced by lower-star filtration of K with respect to -$\rho$
2: Obtain Simplified Discrete Gradient Vector Field
  - Initialize trivial vector field
  - For each persistence pair, perform cancellation if possible and persistence $\leq \tau$
3: Collect Output
  - compute the 1-stable manifold of each critical edge with persistence $> \tau$
4: **return** union of 1-stable manifolds as reconstructed graph

***Step 1.***

Given a 2D-cubic complex K with a density function $\rho : V \to \mathbb{R}$ defined at vertices $V$ of K, we first perform the persistence algorithm to the lower-star filtration of $\rho' = -\rho$. The output is a collection $\Pi_\rho$ of pairs of cells in K (e.g, vertex-edge pairs or edge-square pairs). Note that as mentioned earlier, we use the negation of the density map $\rho' = -\rho$ in our algorithm so that it is easier later to compute the V-path between critical points (minima) and critical edges (index-1 saddles). In our implementation, we adapted DIPHA[40] to compute persistence because it is a distributed persistent homology algorithm which helps to reduce computation time.

***Step 2.***

The second step of the algorithm is to compute and simplify discrete gradient vector field. In theory, one should start with an initial trivial vector field $W$ where there is no discrete gradient vectors (that is, all simplices are critical at the beginning). One can then go through the persistence pairs from the output $\Pi_\rho$ of Step-1: any persistence pair $(\sigma, \tau)$ with persistence $\leq \tau$ is considered as noise, and one then performs the Morse Cancellation operation to cancel them if possible. In the end, this would give rise to a new discrete gradient vector field $W'$, where all low-persistence critical simplices are removed (if possible).

However, to implement this idea, [22] shows that in fact one does not need to explicitly perform Morse cancellations. Instead, all that is needed is to calculate the spanning forest that is made up of all negative edges (edges that are paired with vertices in $\Pi_f$) with persistence less than or equal to $\tau$. Positive edges (edges paired with a square) and edges with persistence greater than $\tau$ are not part of the spanning forest. No explicit discrete gradient vector field needs to be computed nor maintained. This spanning forest contains sufficient information for the Step 3 below. This step takes linear time once the persistence pairings are computed in Step 1.

***Step 3.***

The third step of the algorithm is to compute the 1-stable manifold of each critical edge whose persistence is at least $\tau$ in the simplified discrete gradient vector field. As shown in[22], for each such edge, the 1-stable manifold is equivalent to the union of the edge with the paths from both vertices to the sink of their corresponding tree in the spanning forest computed in Step 2. The union of all 1-stable manifolds is outputted by the algorithm as the reconstructed graph. Again we note that the 1-stable manifolds of -$\rho$ are analogous to the 1-unstable manifolds (mountain ridges) of the density field $\rho$.
We refer the output of this algorithm as the *Morse skeleton graph*, $G$. As shown in Fig. 1, this skeleton graph will then feed to the Simplification Step of the entire pipeline, to further remove noise, false branches and output the final tree/forest summary and produce vectorized objects for quantification.
