## Supplement PSF for "Skeletonization of neuronal processes using Discrete Morse techniques from computational topology"

### S.2 Point Spread Function

In order to estimate an areal line density from the detected fragments, it is necessary to estimate the effective thickness of the optical plane through the sample when acquiring the microscopic image. This thickness is necessarily nonzero due to the finite wavelength of light, and its extent may be estimated by characterizing the point spread function (PSF) of the microscopic imaging systems involved. For the fluorescent Whole Slide Imaging scans using the Nanozoomer HT2.0, we carried out an empirical determination of the PSF, using small fluorescent beads sufficiently smaller than the light wavelength to act as point sources, and also compared with the theoretical PSF. These two estimates sufficiently coincided for our purposes that we adopted the theoretical PSF to estimate the optical plane thickness as detailed below.

#### S.2.1 Theoretical PSF

Various tools exist in the literature for estimating a theoretical PSF for wide field fluorescent microscopy. We used the Richards & Wolf (1959) model as implemented in an open-source package for PSF generation using the Fiji plug-in tool (http://bigwww.epfl.ch/algorithms/psfgenerator/). We used the following input parameters needed in this tool, corresponding to the imaging setup:

- Refractive index: (air) 1
- Wavelength: 660nm
- NA: 0.75
- XY resolution: 460nm
- Z: 130nm

The result theoretical PSF is symmetric to the z=0 plane, forming a cone shape with FWHM (Full Width Half Maximum) of approximately 1.5$\mu m$ in axial (z) direction, as seen in Supplementary Fig. 5.

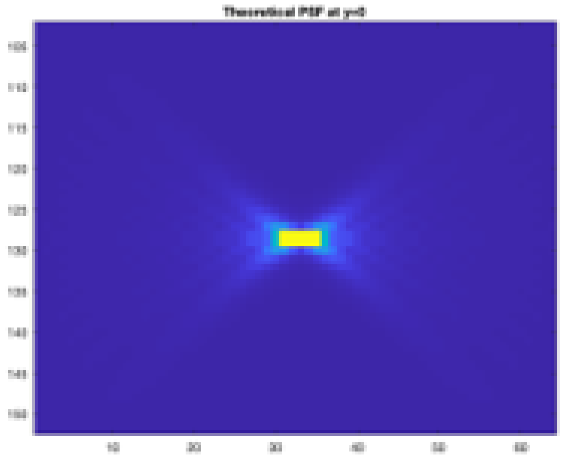

**Supplementary Fig. 5** | Theoretical PSF

#### S.2.2 Experimental PSF

To obtain an empirical PSF of our imaging system, we imaged fluorescent nanobeads of 200nm. By aligning and averaging across 33 samples, we obtained an average experimental PSF as shown in Supplementary Fig. 6.

#### S.2.3 Comparing the theoretical and experimental PSFs

We compared the theoretical and experimental PSFs by normalizing the intensity against the maximum point in the center, and overlaying the two functions within the same coordinate system. The experimental PSF can be seen to correspond approximately to the theoretical PSF in all planes of section as seen by comparing equivalent contour lines (Supplementary Fig. 7). We therefore adopted the FWHM of the theoretical PSF in the estimates in the text. Please note that this provides an overall multiplicative factor to the estimation of line density, so that errors in our estimation would only be reflected in an overall multiplicative factor, and not affect the relative distribution of line density across voxels in the image. For example, it would not affect the surprise estimate which proceeds from normalized probability densities. However, it would impact the estimation of the total length of axons emanating from the injection site. As discussed in the text, this is a biologically meaningful quantity, and therefore work the extra

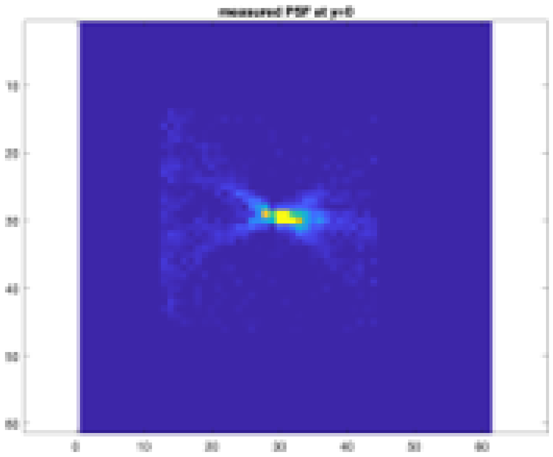

**Supplementary Fig. 6 |** Measured PSF

trouble incurred in trying to evaluate the PSF. The present work should be regarded as a first step in this direction, as there remains room for estimation of the corresponding stereological correction factors.

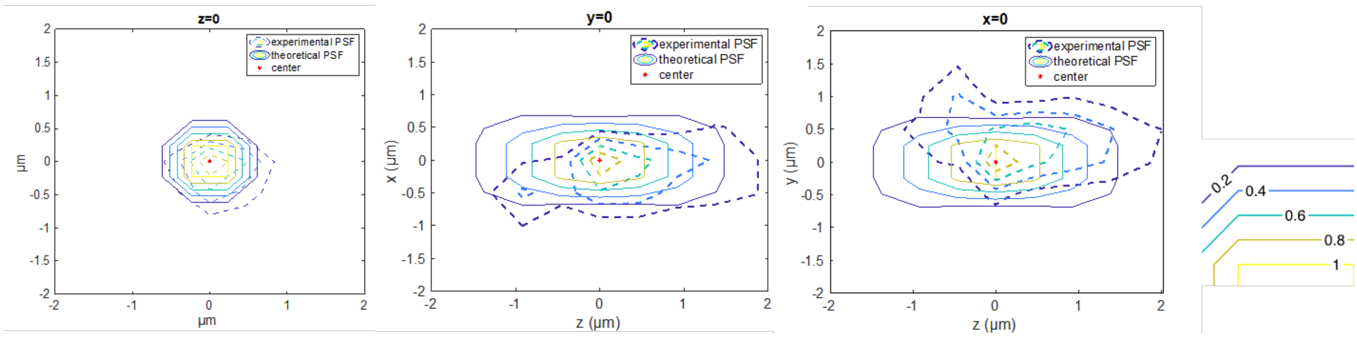


**Supplementary Fig. 7 |** Overlay of experimental and theoretical contours. The same color of the contour lines corresponds to the same level of normalised intensity level, taking the center as 1.
